## Supplementary figures for "Investigating autism associated genes in *C. elegans* reveals candidates with a role in social behaviour"

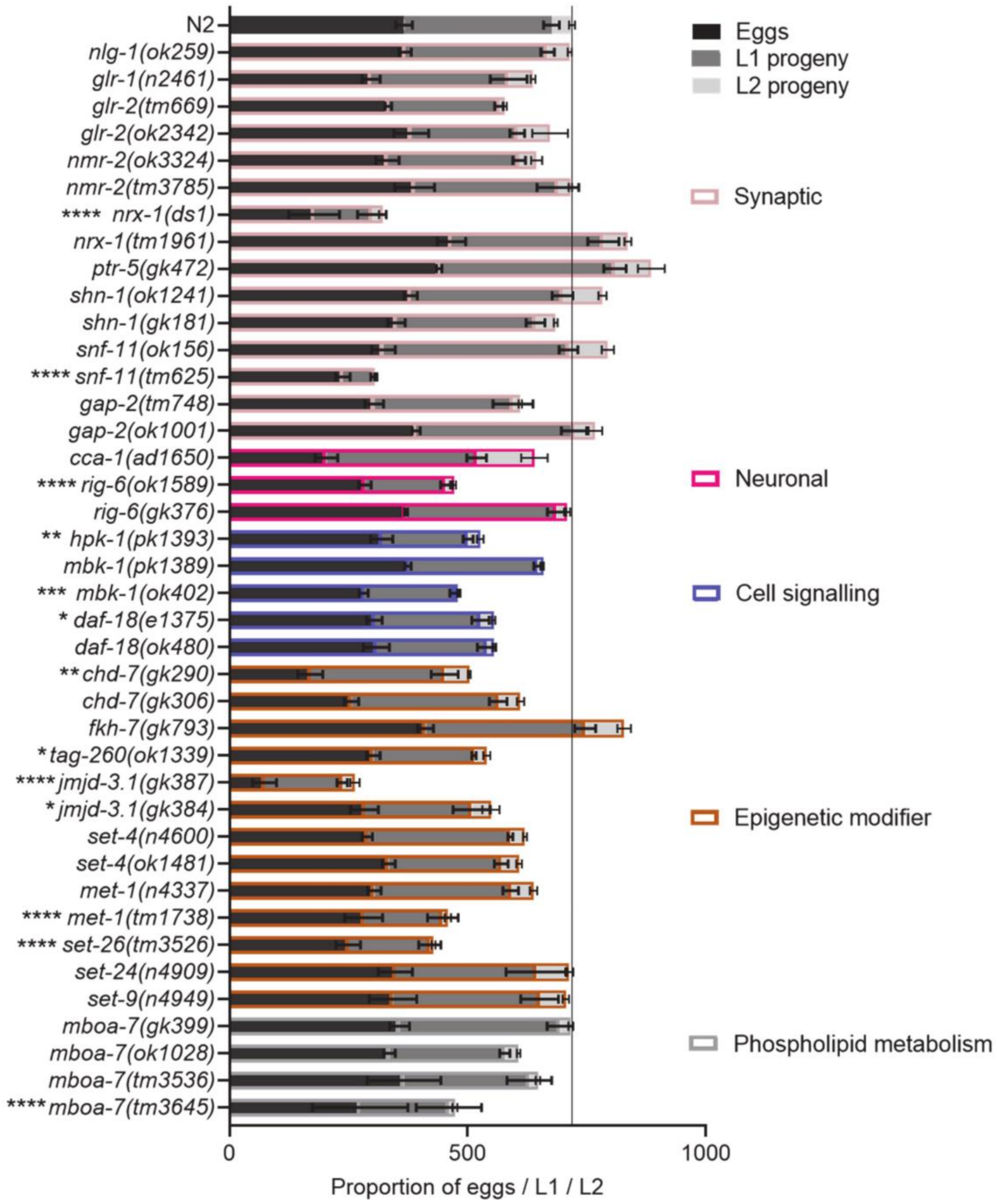


**S1 Fig: Proportion of eggs and progeny produced by *C. elegans* mutants after 24 hours.** After a food leaving assay the number of eggs and progeny produced after 24 hours was counted. N2 and *nlg-1(ok259)* n=19. All other mutants n=3-4. The black line indicates the total number of eggs and progeny produced by N2 control. All data shown as mean ±SEM. Statistical analysis performed using a two-way ANOVA and Dunnetts's multiple comparison test; ns, $p>0.05$; *, $p<0.05$; **, $p\leq0.01$; ***, $p\leq0.001$; ****, $p\leq0.0001$. All significance relates to the total number of eggs and progeny produced in comparison with N2 control.

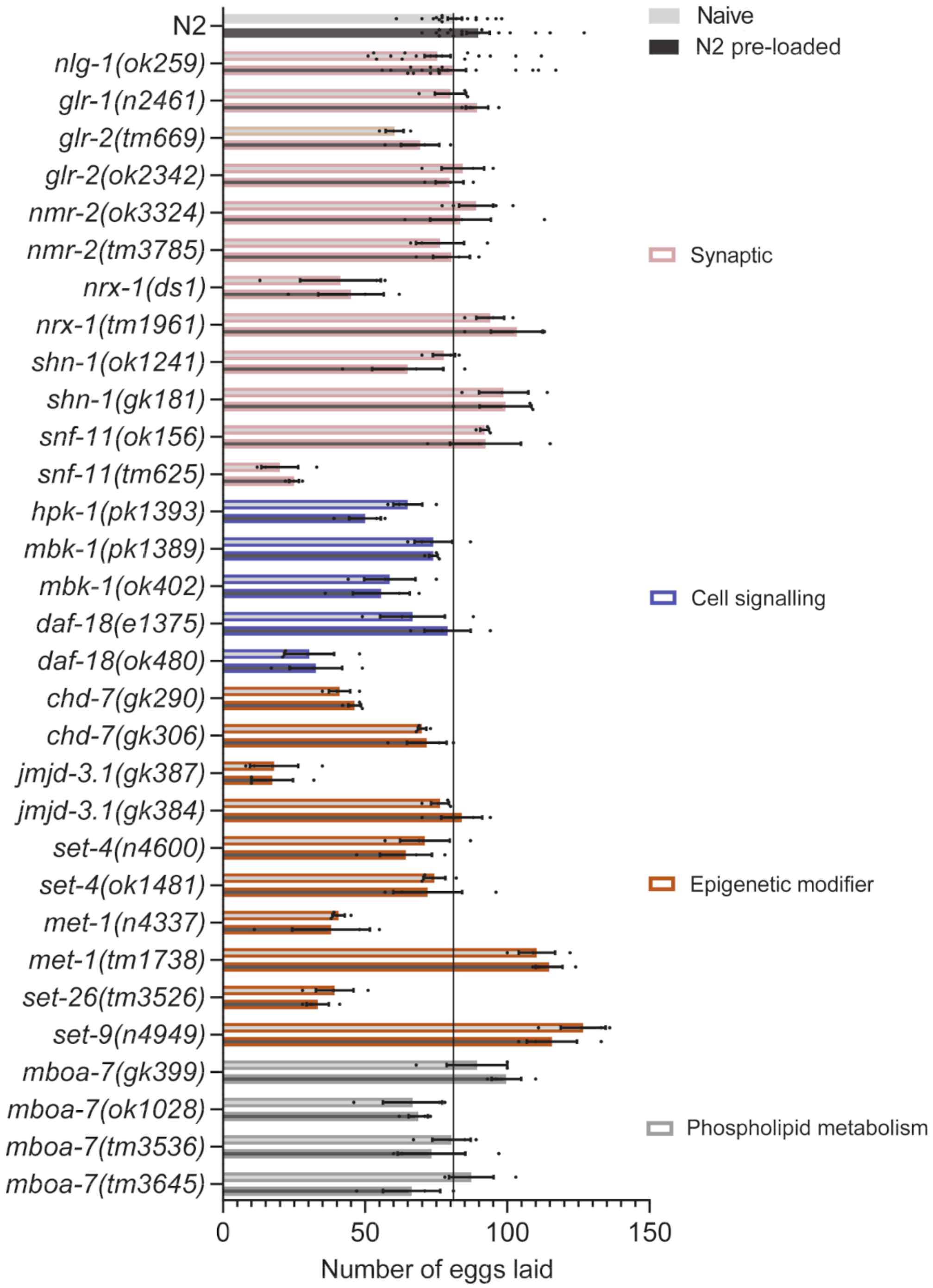


**S2 Fig: The number of eggs laid is unchanged in the presence of progeny.** After a food leaving assay the number of eggs laid was quantified. The black line indicates the number of eggs laid by N2 control. All data shown as mean ±SEM. N2 and *nlg-1(ok259)* n=16. All other mutants n=3-4. Statistical analysis performed using a two-way ANOVA and sidak's multiple comparison test. No significance was identified when comparing eggs laid on a naïve and pre-loaded food lawn for each strain.
